# SLAMF6 deficiency augments tumor killing and skews towards an effector phenotype revealing it as a novel T cell checkpoint

**DOI:** 10.1101/824946

**Authors:** Emma Hajaj, Galit Eisenberg, Shiri Klein, Shoshana Frankenburg, Sharon Merims, Inna Ben-David, Thomas Eisenhaure, Sarah E. Henrickson, Alexandra-Chloé Villani, Nir Hacohen, Nathalie Abudi, Rinat Abramovich, Jonathan Cohen, Tamar Peretz, André Veillette, Michal Lotem

## Abstract

SLAMF6 is a homotypic receptor of the Ig-superfamily whose exact role in immune modulation has remained elusive. Its constitutive expression on resting and activated T cells precludes it from being a *bona fide* exhaustion marker. By breeding Pmel-1 mice with SLAMF6 KO mice, we generated donors for T cells lacking SLAMF6 and expressing a transgenic TCR for gp100-melanoma antigen. Activated Pmel-1xSLAMF6 KO CD8 T cells displayed improved polyfunctionality and strong tumor cytolysis. T-bet was the dominant transcription factor in Pmel-1xSLAMF6 KO cells, and upon activation, they acquired an effector-memory phenotype. Blocking LAG-3 improved the function of SLAMF6 deficient T cells even further. Finally, adoptive transfer of Pmel-1xSLAMF6 KO T cells into melanoma-bearing mice resulted in lasting tumor regression in contrast to temporary responses achieved with Pmel-1 T cells. These results support the notion that SLAMF6 is an inhibitory immune receptor whose absence enables powerful CD8 T cells to eradicate tumors.

## Introduction

The SLAM family of receptors (SFRs) is a set of six receptors expressed on hematopoietic cells (Wu and Veillette, 2016; Cannons, Tangye and Schwartzberg, 2011; Veillette, 2010; Calpe *et al.*, 2008). All SFRs, except 2B4, are homotypic binders, i.e., they engage the same ectodomain sequence, either *in cis* (same cell) or *in trans* (adjacent cell) configuration. Most hematopoietic cell types express 3-5 members of the SLAM family.

SFRs generate signals via a bi-phasic mechanism of recruitment to tyrosines in the immunoreceptor tyrosine-based switch motifs (ITSMs) in their cytoplasmic domain. SLAM associated protein (SAP), a small protein containing the Src homology 2 (SH2)-domain, was shown to be the default adaptor of the SFRs, interchanging with protein tyrosine phosphatases, mainly SHP-1, but also SHP-2, inositol phosphatase SHIP-1 and protein tyrosine kinase Csk (Wu and Veillette, 2016; Cannons, Tangye and Schwartzberg, 2011; Veillette, 2010; Calpe *et al.*, 2008).

SLAMF6, also known as CD352, LY108, or NTB-A, is a homotypic SFR expressed on T cells, NK cells, B cells, and dendritic cells (Bottino *et al.*, 2001; Zhong and Veillette, 2008). Kageyama et al. linked SLAMF6 to the anchoring of T cells to their target cells, and subsequent cytolysis of the target (Kageyama *et al.*, 2012). According to these authors, functional SAP is critical for SLAMF6 activity. In mice lacking SAP, SLAMF6 was shown to inhibit T cell function (Kageyama *et al.*, 2012; Zhao *et al.*, 2012; Bottino *et al.*, 2001). The role of SLAMF6 in healthy T cells expressing normal SAP levels was generally inferred from contextual data and is not yet clear. There are indications that SLAMF6 plays an activating role in double-positive thymocytes (Dutta *et al.*, 2013) along with evidence that it plays an inhibitory role in iNKT cells and CD8 T cells (Lu *et al.*, 2019; Eisenberg *et al.,* 2018). Gene expression profiles of T cell subsets link SLAMF6 to the progenitor-exhausted state (Miller *et al.*, 2019) and to the tuning of the critical number of T cells required for proper differentiation (Polonsky *et al.*, 2018).

To elucidate the net function of SLAMF6, we generated a transgenic mouse with the Pmel-1 melanoma-specific T-cell receptor (TCR) expressed in CD8 T cells, in which the *SLAMF6* gene was knocked out. In this report, we show for the first time that SLAMF6 KO CD8 T cells display improved anti-melanoma activity and prevent melanoma growth more effectively than CD8 T cells with intact and functional SLAMF6. Since SLAMF6 is constitutively expressed on T cells, it acts as an inhibitory checkpoint receptor whose absence allows the eradication of established tumors by CD8 T cells.

## Results

### SLAMF6 is constitutively expressed on T cells and increases upon activation

SLAMF6 is an immune receptor constitutively expressed on non-activated and activated T cells (Eisenberg *et al.*, 2018). The level of SLAMF6 transcription and receptor expression, however, is dynamic, changing with time and activation states. To record SLAMF6 expression in a longitudinal manner, human tumor-infiltrating lymphocytes (TILs) were activated for five days, and SLAMF6 transcript and protein expression were measured (Figures 1A-C). After one day of activation, there was an initial decrease in the SLAMF6 transcript that switched to over-expression (Figure 1C). From three days after activation onward, SLAMF6 receptor expression consistently increased (Figures 1A and B). Interestingly, the increased expression was most pronounced in T cells activated in the absence of IL-2 (Figure 1D). A similar pattern was observed for the expression of the murine SLAMF6 receptor on Pmel-1 CD8 T cells (Figure 1E).

**Figure 1:**
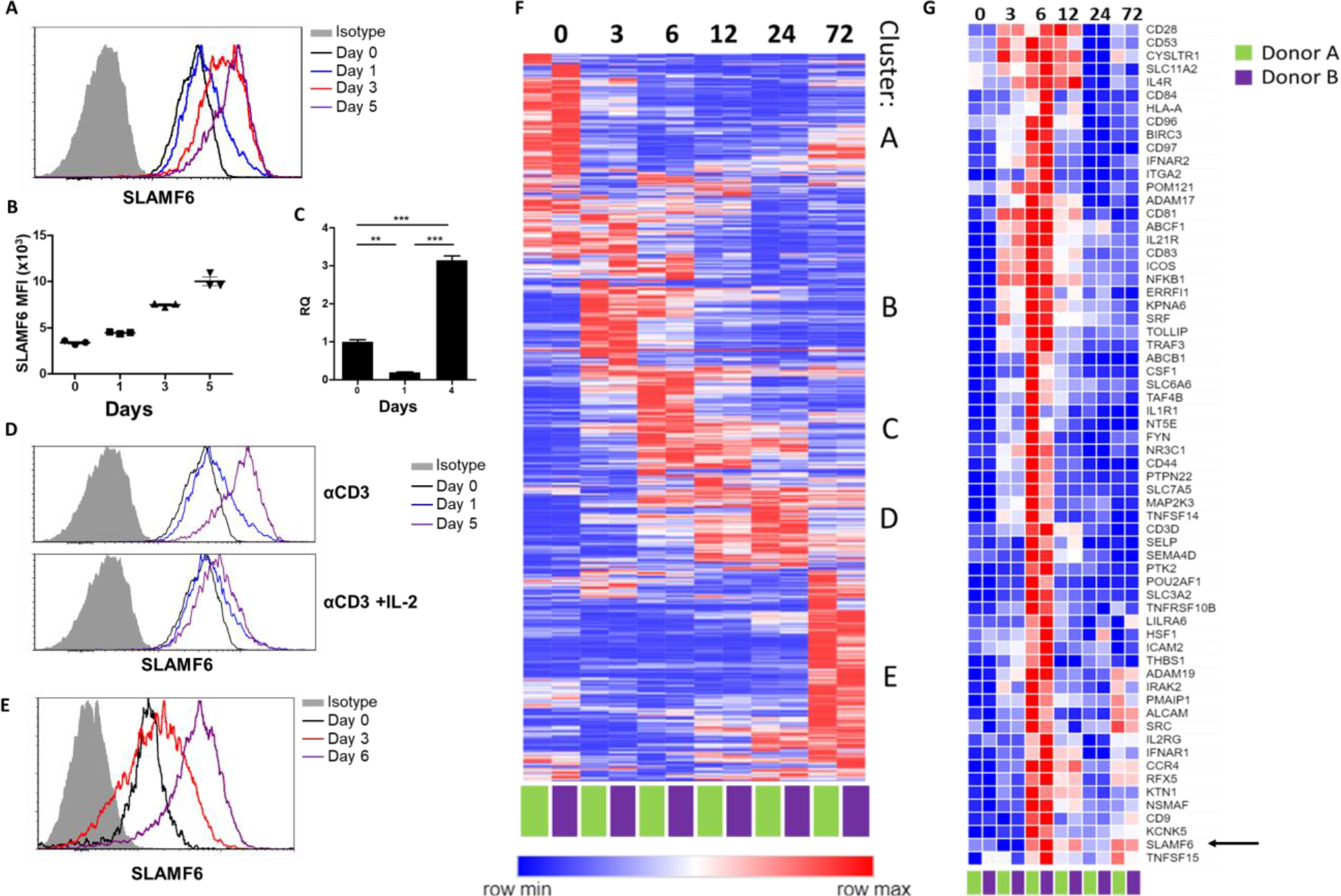
SLAMF6 is constitutively expressed on T cells and increases upon activation. **A-C**, SLAMF6 expression in human TIL412 cells, activated for five days. **A**, flow cytometry at the indicated time points. **B**, Median fluorescence intensity (MFI) of SLAMF6, days 1-5. **C**, quantitative RT-PCR for *SLAMF6*. RNA was extracted at the indicated time points. Data normalized to *HPRT* expression at each time point and to the basal expression level on day 0. One-way ANOVA. **, P < 0.01, ***, P < 0.001. **D**, SLAMF6 expression by flow cytometry in human TIL412 cells activated for 5 days with anti-CD3 or with anti-CD3 plus IL-2, at the indicated time points. **E**, SLAMF6 expression by flow cytometry in Pmel-1 mouse splenocytes activated for 6 days, at the indicated time points. **F**, Row normalized expression of immune-related genes from RNAseq, clustered according to similar expression patterns. CD4+ T cells from two donors were stimulated with anti-CD3 plus anti-CD28 for 72 hours, RNA was extracted and sequenced. Numbers in the top panel indicate hours. **G,** Magnification of cluster C. *SLAMF6* is marked.

To identify other immune-related genes that may cluster with SLAMF6, longitudinal RNA sequencing data were generated from CD4 T cells from two healthy human donors. Five groups of genes (clusters A-E) were identified (Figure 1F). Cluster A represents genes highly expressed in non-activated cells, and downregulated upon activation, such as *BCL2* and *JAK1*. Cluster B represents fast-rising genes undergoing transcription as early as 3-6 hours following activation, but down-regulated after that; genes in this cluster include the activation marker *CD69*. Clusters C and D include intermediate genes, upregulated after 6 hours (C) or between 12-24 hours (D), and downregulated later. Lastly, cluster E includes late-rising genes, such as *LAG3*. The *SLAMF6* transcript appears in cluster C, rising at 6 hours of activation and staying high after that (Figure 1G). Other genes in cluster C are *CD44*, encoding a glycoprotein that takes part in T cell activation (Huet *et al.*, 1989), and *CD28* and *ICOS*, which encode co-stimulatory immune receptors (Turka *et al.*, 1990; Dong, Juedes and Temann, 2001). The increase in the transcription of receptors that are stably expressed at all times may hint at enhanced recruitment and degradation of these receptors during activation, possibly in the immune synapse (Onnis and Baldari, 2019).

### SLAMF6 expressed *in trans* by a melanoma target inhibits anti-tumor T cell reactivity

Because SLAMF6 is constantly present on T cells, it is difficult to decipher its effect when it acts as a ligand, introduced *in trans*. To solve this problem, we generated a cell line derived from B16-F10/mhgp100 melanoma, over-expressing SLAMF6 (Figure 2A). Compared to the wild-type cell line, the SLAMF6-expressing melanoma cells co-cultured with Pmel-1 CD8 T cells led to decreased IFN-γ secretion by the lymphocytes (Figures 2B and C). To evaluate the effect of SLAMF6 expression on melanoma rejection *in vivo*, SLAMF6-expressing melanoma was compared to the parental B16-F10/mhgp100 line, in an adoptive T cell transfer regimen, using activated Pmel-1 CD8 T cells (Figure 2D). The B16-F10/mhgp100/SLAMF6^+^ tumors grew more aggressively: on day 23, the mean tumor volume was 380 mm^3^ in the SLAMF6-expressing melanomas, compared to 137 mm^3^ in the non-modified tumors (Figure 2E).

**Figure 2:**
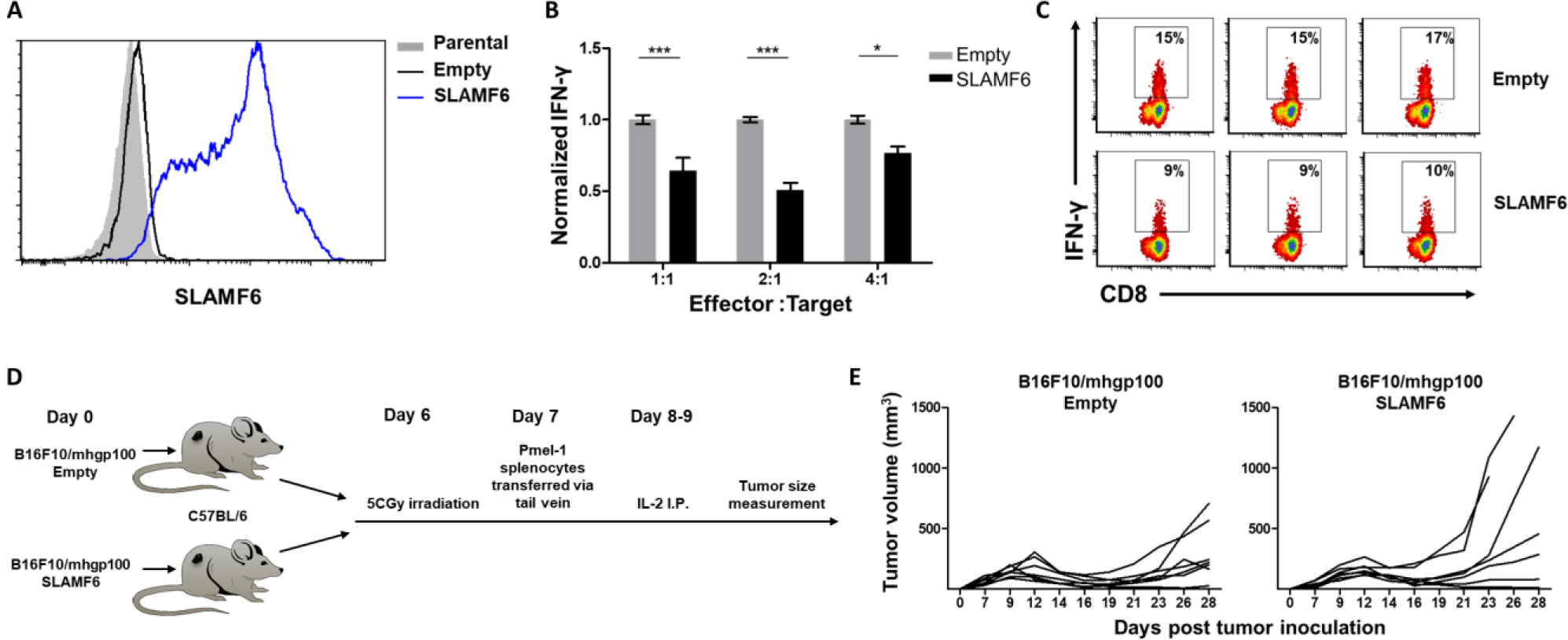
SLAMF6 expressed *in trans* by a melanoma target inhibits anti-tumor T cell reactivity. **A**, SLAMF6 expression on B16-F10/mhgp100 parental or transfected (SLAMF6 or empty) melanoma cells. **B**, Pmel-1 splenocytes were activated for 7 days with gp100_25-33_ peptide and IL-2 (30 IU/ml), and then incubated overnight with B16-F10/mhgp100/empty or B16-F10/mhgp100/SLAMF6 melanoma cells at the indicated effector-to-target ratios. IFN-γ secretion was measured by ELISA. **C**, Pmel-1 splenocytes were activated for 7 days with gp100_25-33_ peptide and IL-2 (30 IU/ml), and then incubated overnight with B16-F10/mhgp100/empty or B16-F10/mhgp100/SLAMF6 melanoma cells. IFN-γ production was detected by intracellular staining and flow cytometry (gated on CD8). Three replicates. **D, E**, Pmel-1 splenocytes were expanded with gp100_25-33_ peptide (1μg/ml) and IL-2 (30 IU/ml) for 7 days. On day 7, cells were transferred i.v. into irradiated C57Bl/6 mice bearing palpable (1 week) B16-F10/mhgp100/empty or B16-F10/mhgp100/SLAMF6 tumors. IL-2 (0.25×10^6^ IU) was administered i.p. twice a day for two days. Tumor growth was measured twice a week. Mice were sacrificed when the tumor reached 15mm in diameter. **D**, Scheme showing experimental layout. **E**, Spider plot showing tumor volume [calculated as L (length) x W (width)^2^ x 0.5]. One-way ANOVA. *, P < 0.05, **, P < 0.01, ***, P < 0.001.

These experiments show that trans-activation of SLAMF6 on lymphocytes, which in this system was achieved with the SLAMF6-expressing melanoma, inhibits the melanoma-specific CD8 T cell response and allows rapid tumor growth.

### Establishment of Pmel-1 x SLAMF6 KO mice as a source of SLAMF6-KO antigen-specific lymphocytes

To evaluate the role of SLAMF6 in melanoma-cognate T cells, we generated a new mouse strain by breeding Pmel-1 mice with SLAMF6 KO mice. The offspring of this cross represented a new strain, which could serve as a source of CD8 T cells lacking SLAMF6 and expressing the transgenic TCR against the H-2D^b^ gp100: 25-33 peptide (Figure 3A). Evaluation of the lymphocyte subsets in these mice showed a lower percentage (15%) of CD8 cells in Pmel-1 x SLAMF6 KO mouse spleens compared to Pmel-1 splenocytes (24%) (Figure 3B and Supplementary Figure 2A). Despite the smaller percent of CD8 cells in the spleens of the SLAMF6-deficient mice, the ratio of CD8 subpopulations (naïve, effector, effector memory, and central memory) was similar in both mouse strains (Figure 3C).

**Figure 3:**
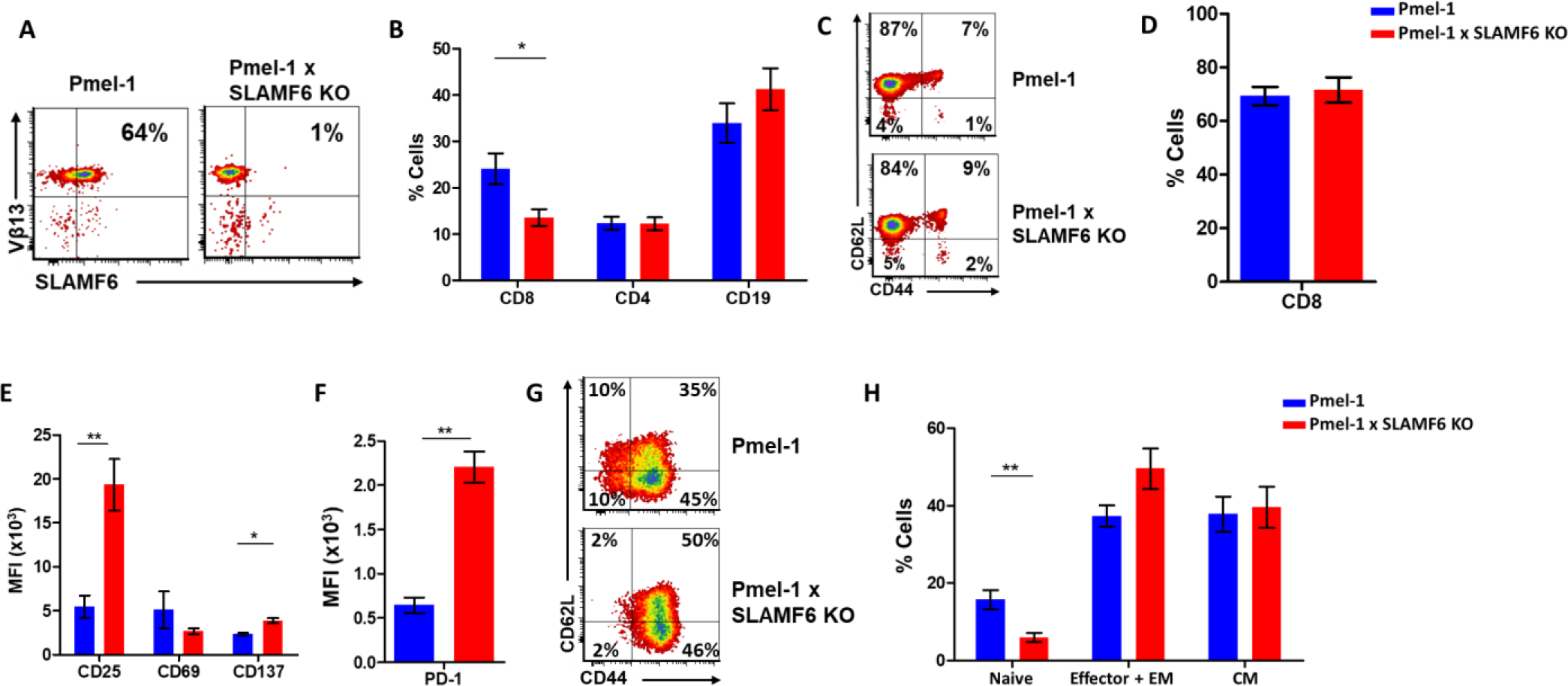
Establishment of Pmel-1 x SLAMF6 KO mice as a source of SLAMF6-KO antigen-specific lymphocytes. **A**, SLAMF6 and Vβ13 expression in Pmel-1 or Pmel-1 x SLAMF6 KO splenocytes measured by flow cytometry. **B**, Percent CD8, CD4, and CD19 cells in spleens from Pmel-1 or Pmel-1 x SLAMF6 KO untreated mice. **C**, Pmel-1, and Pmel-1 x SLAMF6 KO CD8 untreated splenocytes were stained with anti-CD44 and anti-CD62L. One representative experiment is shown. **D**, Percent CD8 cells in Pmel-1 or Pmel-1 x SLAMF6 KO splenocytes after 7 days of *in vitro* activation with gp100_25-33_ peptide and IL-2 (30 IU/ml). **E**, Flow cytometry for activation markers (CD25, CD69, CD137) in Pmel-1 or Pmel-1 x SLAMF6 KO splenocytes after 3 days of *in vitro* activation, as in (D). Median fluorescence intensity (MFI) is shown. **F**, Expression of PD-1 in Pmel-1 or Pmel-1 x SLAMF6 KO CD8 T cells after 7 days of *in vitro* activation, as in (D). Median fluorescence intensity (MFI) is shown. **G, H**, After 7 days of activation, Pmel-1 and Pmel-1 x SLAMF6 KO CD8 T cells were stained with anti-CD44 and anti-CD62L. CD8 subpopulations were defined for each mouse strain. **G**, One representative experiment and **H**, summary of subpopulations identified by flow cytometry in 5 experiments is shown. EM, effector memory, CM, central memory. Student t-test. *, P < 0.05, **, P < 0.01, ***, P < 0.001.

In the initial *in vitro* activation assays, it was already clear that the Pmel-1 x SLAMF6 KO T cells have improved functional capacity. Their proliferative response to peptide stimulation was preserved, which was mandatory to produce ample numbers of CD8 lymphocytes for the adoptive T cell transfer regimen (Figure 3D). CFSE dilution curves were identical in the two mouse strains, as were the activation-induced cell death (AICD) rates (Supplementary Figures 2B and C). After three days of activation, the KO mice had higher expression levels of CD25 and CD137 (4-1BB) activation markers (Figure 3E). In parallel, higher PD-1 expression was detected on day 7 (Figure 3F), which in this experimental context was initially taken as an indicator of activation, but was later attributed to SAP deficiency in the SLAMF6 KO lymphocytes (see below in Results and Figures 5A and B). PD-1 overexpression was also noted by Lu et al. in iNKT cells from SFR KO mice (Lu *et al.*, 2019). The expression of other SLAM family members and ligands (CD48, LY9, CD244, CD84, CD319) on T cells during activation was similar in Pmel-1 and Pmel-1 x SLAMF6 KO cells (Supplementary Figure 2D).

Lastly, we phenotyped the T cells following 3-day and 7-day activation to assess subset ratios based on CD44 and CD62L differentiation markers. While initially, all Pmel-1 T cells were naïve, only a negligible number of the activated Pmel-1 x SLAMF6 KO cells remained in the naïve CD62L^high^/CD44^low^ state compared to 10% of the Pmel-1 cells. The complete shift of the activated Pmel-1 x SLAMF6 KO lymphocytes towards effector and effector memory phenotypes indicates the strength of their response to activation (Figures 3G and H).

Pmel-1 x SLAMF6 KO lymphocytes, generated to evaluate the effect of SLAMF6 on antigen-specific activated T cells, showed stronger activation and global acquisition of an effector phenotype. We attribute these results, in contrast to those previously obtained with lymphocytes from SLAMF6 KO mice, to the role of SLAMF6 in the immune synapse, as the results could only be obtained if a synapse formation had been initiated via cognate TCRs.

### Pmel-1 x SLAMF6 KO T cells have a better functional capacity

In the previous experiments, the effect of SLAMF6 deletion was evaluated in the activation and proliferation phases. To test if this superior activation also affects anti-tumor immunity, we characterized Pmel-1 x SLAMF6 KO melanoma-specific T cells in the effector phase.

Comparing IFN-γ secretion in response to melanoma cells by Pmel-1 versus Pmel-1 x SLAMF6 KO lymphocytes showed that cytokine production by the Pmel-1 x SLAMF6 KO lymphocytes was significantly higher at all effector-to-target ratios (p=0.05, Figures 4A and B and Supplementary Figures 3A and B). In addition to IFN-γ, higher secretion of GM-CSF and lower levels of IL-10 and IL-13 were measured in the SLAMF6 KO T cells (Figures 4C and D). Since GM-CSF is a strong recruiter of innate immune cells, while IL-10 and IL-13 drive suppressor traits, this secretion profile supports autocrine and paracrine immune activation. mRNA data validated the secretion assays (Supplementary Figure 3C). Importantly, Pmel-1 x SLAMF6 KO T cells produced higher levels of granzyme B in response to melanoma compared to Pmel-1 T cells, which are already strong killers due to their TCR design (Figures 4E and F). These results indicate that in the absence of the SLAMF6 modulatory effect, even strong cytolysis can be further enhanced.

**Figure 4:**
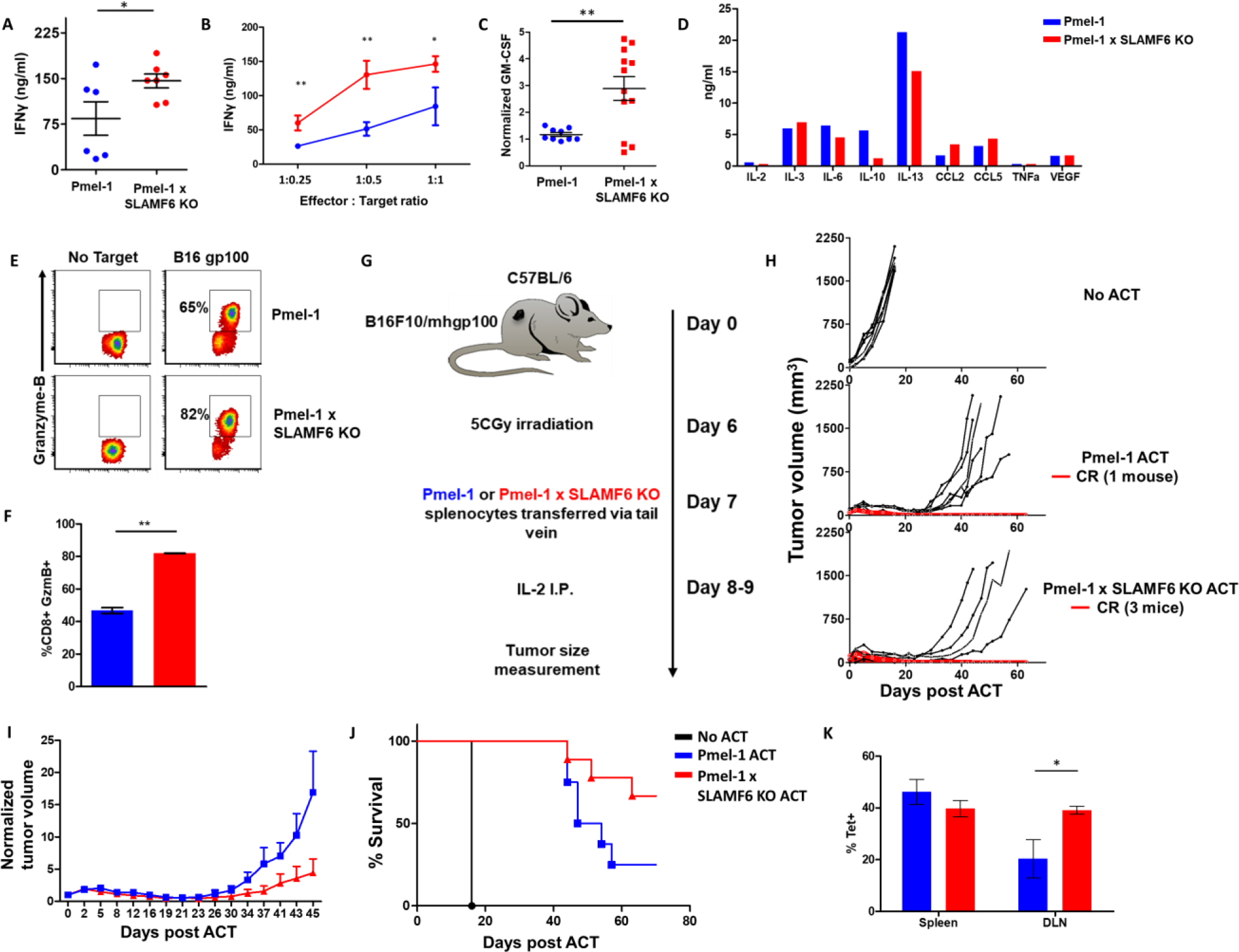
Pmel-1 x SLAMF6 KO T cells have a better functional capacity. **A-D**, Pmel-1 or Pmel-1 x SLAMF6 KO splenocytes were activated for 7 days with gp100_25-33_ peptide and IL-2 (30 IU/ml) and then incubated overnight with B16-F10/mhgp100 melanoma cells. **A,** The cells were incubated at a 1:1 effector-to-target ratio. IFN-γ secretion was measured by ELISA. Each point represents one mouse. **B,** The cells were incubated at the indicated effector-to-target ratios. IFN-γ secretion was measured by ELISA. **C,** The cells were incubated at a 1:1 effector-to-target ratio. GM-CSF secretion was measured by ELISA. Each point represents one mouse. **D**, Conditioned medium was collected and analyzed with Quantibody mouse cytokine array. **E, F**, Pmel-1 or Pmel-1 x SLAMF6 KO splenocytes were activated for 7 days with gp100_25-33_ peptide and IL-2 (30 IU/ml) and then incubated for 16 hours with B16-F10/mhgp100 melanoma cells. Granzyme-B expression was detected by flow cytometry. One representative experiment (**E)**, and a summary of triplicates (**F)**, are shown. **G-J**, B16-F10/mhgp100 mouse melanoma cells were injected s.c. into the back of C57BL/6 mice. Pmel-1 or Pmel-1 x SLAMF6 KO mouse splenocytes were expanded with gp100_25-33_ peptide in the presence of IL-2 (30 IU/ml). On day 7, Pmel-1 cells or Pmel-1 x SLAMF6 KO cells were adoptively transferred i.v. into the irradiated tumor-bearing mice. N=8 mice per group. Tumor size was measured 3 times a week. **G**, Scheme of the experimental layout. **H**, Spider plots showing tumor volume [calculated as L (length) x W (width)^2^ x 0.5]. CR, complete response. **I,** Normalized tumor volume (Mean±SEM) until day 45, on which the first mouse had to be sacrificed. Tumor dimensions were normalized to the 1^st^ measurement. **J**, Kaplan Meier survival curve. **K**, Percent T cells specific for gp100_25-33_ peptide in the spleen or tumor draining lymph nodes (DLN) of mice sacrificed 7 days post ACT. Tet, tetramer. Student t test. *, P < 0.05, **, P < 0.01, ***, P < 0.001.

To evaluate the anti-tumor activity of Pmel-1 x SLAMF6 KO cells, we assessed adoptive cell transfer (ACT) of 7-day pre-activated gp100:25-33-specific, Pmel-1 or Pmel-1 x SLAMF6 KO CD8 T cells, transferred into mice bearing palpable B16-F10/mhgp100 melanoma in their back skin, followed by a 2-day course of intraperitoneal IL-2 (Figures 4G-J). The spider curves comparing melanoma growth following Pmel-1 versus Pmel-1 x SLAMF6 KO T cell transfer revealed that in the first four weeks post-transfer, tumor growth was inhibited in both groups. However, on week four, tumors treated with Pmel-1 ACT escaped control and grew again in six of the seven animals, whereas mice receiving Pmel-1 x SLAMF6 KO ACT survived longer, and three of the seven treated mice remained tumor-free for over 80 days (Figure 4H). Vitiligo was noted in all mice attaining complete response, showing up earlier in the KO group, at the 6^th^ week, indicating the strength of the response (Supplementary Figure 3D).

In a similar experiment, peptide-activated Pmel-1 CD8 T cells or Pmel-1 x SLAMF6 KO CD8 T cells were transferred into mice bearing 7-day-old tumors. A week later, mice were sacrificed, and spleens, tumors, and tumor-draining lymph nodes were extracted and evaluated for the presence of transferred T cells, using flow cytometry to detect the gp100_25-33_ tetramer. A higher proportion of gp100_25-33_ tetramer^+^ cells was found in the draining lymph nodes of mice that had received Pmel-1 x SLAMF6 KO cells (Figure 4K) compared to those who had received Pmel-1 cells. Tumors from both groups had a similar density of infiltrating lymphocytes (Supplementary Figure 3E).

In summary, melanoma-specific T cells lacking SLAMF6 showed improved functional capacity both *in vitro* and *in vivo* and induced longer-lasting tumor remission with longer tumor control compared to their wild type counterparts.

### Mechanism associated with the inhibitory function of SLAMF6

The goal of the next series of experiments was to identify mechanisms underlying the improved effector function of Pmel-1 x SLAMF6 KO lymphocytes. We initially evaluated the level of SAP, the primary adaptor required for SLAMF6 signaling, encoded by the *Sh2d1a* gene, which was intact in the SLAMF6 KO mice (Figure 5A). SAP is a critical adaptor that recruits Fyn kinase to SLAMF6. However, while SAP transcript was found at similar levels in WT and SLAMF6 KO lymphocytes, SAP protein was not detectable in SLAMF6 KO cells (Figure 5B). The discrepancy between the transcript and the protein can be explained by rapid degradation of cytoplasmic SAP in its unbound form. SAP protein deficiency also implies that SLAMF6 is its major anchor in non-activated CD8 T cells, despite the fact that SAP also mediates the inhibitory activity of PD-1 (Peled *et al.,* 2018). As shown in Figure 3F, PD-1 expression is more than two-fold increased in 7-day activated Pmel-1 x SLAMF6 KO lymphocytes compared to Pmel-1, but, as we have shown, this gap did not affect functionality. Thus, PD-1 overexpression can represent the failure of this receptor to generate a negative feedback loop in over-activated T cells.

**Figure 5:**
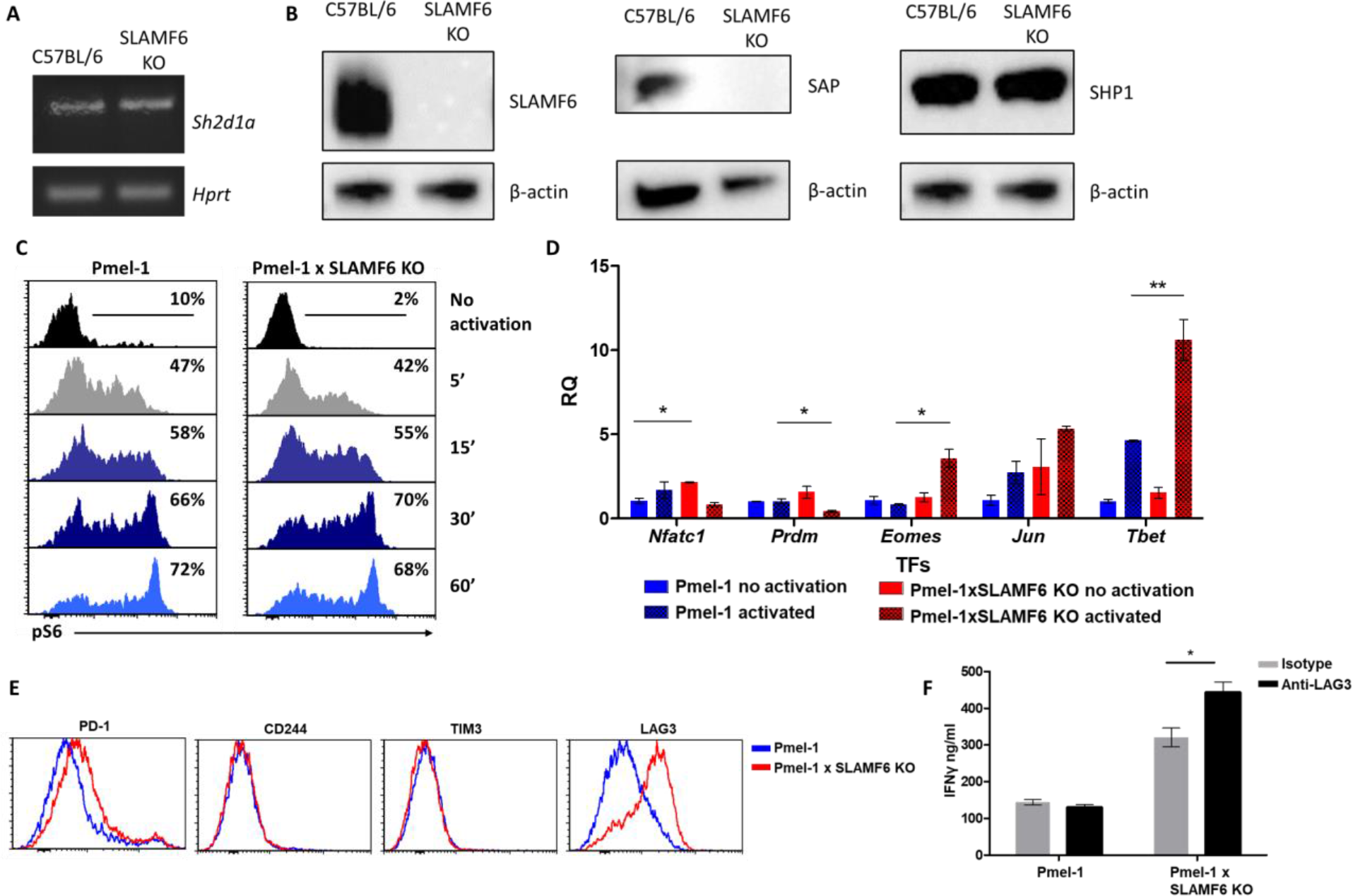
Mechanism associated with the inhibitory function of SLAMF6. **A**, RNA expression of *Sh2d1a* transcript (SAP) in WT and SLAMF6 KO splenocytes. **B,** Immunoblot analysis of expression of SLAMF6, SAP and SHP-1 in WT and SLAMF6 KO splenocytes. **C,** Pmel-1 and Pmel-1 x SLAMF6 KO splenocytes were activated with gp100_25-33_ peptide for the indicated time points. At the end of the activation, cells were fixed and stained for phosphorylated S6. **D,** Pmel-1 and Pmel-1 x SLAMF6 KO splenocytes were either activated with gp100_25-33_ peptide in the presence of IL-2 (30 IU/ml) for 18 hours or kept only with IL-2 for 18 hours (non-activated). After 18 hours, the cells were lysed, RNA was extracted, and quantitative RT-PCR for transcription factors expression was performed. Data was normalized to *Hprt* expression for each mouse strain. Values for each condition were normalized to Pmel-1 non-activated values for each gene. **E**, Pmel-1 and Pmel-1 x SLAMF6 KO splenocytes were expanded with gp100_25-33_ peptide in the presence of IL-2 (30 IU/ml) for 7 days. After the expansion phase, the cells were kept for an additional 5 days without supplements. Expression of exhaustion markers was measured in Pmel-1 or Pmel-1 x SLAMF6 KO splenocytes. **F**, Pmel-1 and Pmel-1 x SLAMF6 KO splenocytes were expanded with gp100_25-33_ peptide in the presence of IL-2 (30 IU/ml) and 10μg/ml anti-LAG-3 or isotype control for 7 days, and then incubated overnight with B16-F10/mhgp100 melanoma cells at a 1:1 effector-to-target ratio. IFN-γ secretion was measured by ELISA. *, P < 0.05, **, P < 0.01.

Next, we measured by flow cytometry the level of phosphorylated ribosomal protein S6 (rpS6), an integrator of important signaling pathways, including PI3K/AKT/mTOR and RAS-ERK (Figure 5C). No difference in phosphorylated rpS6 was found in the Pmel-1 x SLAMF6 KO T cells.

Therefore, we proceeded to identify transcription regulators whose activity differed between the Pmel-1 cells and the Pmel-1 x SLAMF6 KO cells. The most prominent regulator found was T-bet, which increased more than twofold in activated Pmel-1 x SLAMF6 KO splenocytes, followed by Eomes (Figure 5D and Supplementary Figure 4). T-bet was originally described as the key transcription factor defining type 1 T helper (Th) cells; it has since been found to play a major role in the acquisition of effector functions by CD8 T cells (Szabo *et al.*, 2000). Type 1 inflammatory signals induce T-bet expression (Joshi *et al.*, 2007), which is in line with the intensive IFN-γ secretion we observed by the Pmel-1 x SLAMF6 KO lymphocytes (Figure 4A).

Lastly, the level of immune receptors that mediate exhaustion was recorded during prolonged activation. Five days after the end of a seven-days activation course, Pmel-1 x SLAMF6 KO T cells displayed similar levels of PD-1, CD244 (SLAMF4), and TIM-3 to those in the Pmel-1 cells, but higher expression of LAG-3 (Figure 5E). Hypothesizing that LAG-3 represents a compensatory mechanism for the enhanced activation of Pmel-1 x SLAMF6 KO T cells, we used a blocking antibody against LAG-3 in the gp100 activation assay. We then measured IFN-γ secretion by activated T cells in response to B16-F10/mhgp100 melanoma cells. As expected, blocking LAG-3 on Pmel-1 x SLAMF6 KO lymphocytes increased their cytokine secretion significantly, whereas it did not affect Pmel-1 cells. Overall, knocking out SLAMF6 together with LAG-3 blockade resulted in a three-fold increase in IFN-γ production (Figure 5F).

## Discussion

The aim of this study was to characterize the role of SLAMF6 in CD8 T cells, in the context of an anti-tumor response. The data obtained identify SLAMF6 as a receptor whose absence significantly improves CD8-mediated tumor regression, suggesting that it is an inhibitory checkpoint.

Historically, SFRs were studied for their part in X-linked lymphoproliferative disease (XLP), a complex genetic immune dysfunction caused by a SAP mutation. XLP is characterized by a compromised immune response to Epstein-Barr virus (EBV) but also by unrestrained T lymphoblast proliferation, which is not necessarily EBV-induced. Thus, it is unclear whether loss of SAP converts all SFRs into “super-inhibitory” receptors or whether, on the contrary, loss of SAP unleashes lymphocytes to proliferate, free from re-stimulation-induced apoptosis (Katz *et al.*, 2014; Kageyama *et al.*, 2012; Zhao *et al.*, 2012; Bottino *et al.*, 2001). Since SAP is an adaptor common to all SLAM family receptors, the role of each individual receptor was obscured by the shared defect. In this situation, SLAMF6 was considered a receptor with a dual function, depending on the interplay between SAP and SHP-1 and SHP-2, protein phosphatases that bind to tyrosines on the cytoplasmic tail of the receptor (Veillette, 2010; Cannons, Tangye and Schwartzberg, 2011; Detre, Keszei and Romero, 2010). SLAMF6 duality was echoed in data from Veillette that showed differing effects of SLAMF6 on NK cells, enhancing function in the priming phase while suppressing cells in the effector-phase (Wu *et al.*, 2016). Also, mice lacking individual SFRs exhibit minor immune deviations (Wu and Veillette, 2016; Cannons, Tangye and Schwartzberg, 2011; Veillette, 2010; Calpe *et al.*, 2008).

In the past, we described that targeting SLAMF6 with its soluble ectodomain yielded CD8 T cells that do not need IL-2 supplementation, either *in vitro* or *in vivo,* to eradicate established melanoma (Eisenberg *et al.*, 2018). The beneficial effect of the soluble ectodomain of SLAMF6 prompted us to generate melanoma-specific SLAMF6 KO T cells, to characterize the role of the receptor in a solid tumor model.

A key finding using the new Pmel-1 x SLAMF6 KO mice described in this manuscript is the absence, in fact, of a dichotomy in SLAMF6 action in effector T cells. On the contrary, knocking-out SLAMF6 in murine antigen-specific CD8 T cells disclosed an unequivocal inhibitory role for the receptor. In its absence, TCR triggering of anti-melanoma CD8 T cells yielded a strong effector phenotype, higher IFN-γ secretion, improved cytolysis, and better outcomes in the adoptive transfer of SLAMF6 KO anti-melanoma CD8 T cells to treat established melanoma. This study identifies SLAMF6 as a powerful inhibitor of anti-tumor immune response. The absence of viable SAP in SLAMF6 KO lymphocytes hints that this adaptor takes a major part in the inhibitory effect of SLAMF6.

To explore the role of SLAMF6 in T cells without the confounding effects of its function in other cell types, we generated a system in which effector T cells interact with their tumor target based on specific epitope recognition and subsequently generate an immunological synapse. The synapse is a subcellular structure involved in the effect of SLAMF6 and is crucial for its study (Zhao *et al.*, 2012). However, although we revealed the inhibitory effect of SLAMF6 in the Pmel-1 x SLAMF6 KO mice, the source and configuration of SLAMF6/SLAMF6 homotypic binding in the wild-type situation were still difficult to characterize. We had to generate a SLAMF6-positive B16-F10/mhgp100 melanoma line to measure the effect, or more exactly, the degree of suppression, that SLAMF6 trans-activation has on the capacity of melanoma-cognate CD8 T cells to eradicate tumors. As shown (Figure 2E), the SLAMF6-expressing melanoma suppressed T cell efficacy and consequently had accelerated growth. Thus, SLAMF6 trans-activation is suppressive in the response of CD8 T cells to melanoma. In experiments in which the only sources of SLAMF6 ligation were fraternal T cells, we cannot rule out *in cis* interactions, i.e., homotypic binding of SLAMF6 molecules on the same cell.

The molecular mechanisms underlying the increased functional capacity of Pmel-1 T cells lacking SLAMF6 have common features with XLP, as the absence of SAP implies. But while XLP is a global defect of all cell types of the immune system, and therefore yields mixed derangements, the absence of SLAMF6 is remarkable for the enhanced functionality of CD8 T cells, in which it is the dominating SFR.

The transcriptional landscape of SLAMF6 KO T cells was governed by the higher expression of *T-bet*. T-bet is a transcription factor that contributes to Th1 and Th17 phenotypes in CD4 T cells. T-bet is prevalent in cytolytic innate lymphocytes residing in tissues and B cells of mouse strains prone to autoimmunity (Plank *et al.*, 2017; Nixon and Li, 2018). The increased activity of T-bet in SLAMF6 KO CD8 T cells implies that T-bet-regulated pathways may operate in CD8 T cells in the absence of functioning SLAMF6, generating ‘type 1’ inflammatory traits and high cytotoxicity. The improved production of IFN-γ and GM-CSF, in parallel with reduced IL-10 and IL-13, is also typical for type 1 phenotypes.

SLAMF6 should be distinguished from typical exhaustion markers because it is expressed on CD8 T cells, regardless of their state of activation. Yigit et al. suggested that blocking SLAMF6 using an antibody can correct the exhaustion phase of T cells (Yigit *et al.*, 2019), but we favor the notion that SLAMF6 hampers T cells at any stage, as reflected from the functional superiority of short-term activated Pmel-1 T cells. Depleting SLAMF6 improved CD8 T cells in the short and long-term, as was most evident when the WT Pmel-1 cells induced the regression of melanoma only for a limited period while the Pmel-1 x SLAMF6 KO cells led to lasting responses in mice (Figure 4H).

While searching for new immunotherapeutic targets, the field of immunotherapy is moving to combination therapies, and to biomarker-based treatment choices, to target the escape mechanisms used by tumors. From the results presented here, we conclude that SLAMF6 is an important checkpoint receptor with a significant inhibitory effect on T cells. The balance between SLAMF6 and LAG-3 and the reciprocal rise of the latter suggests that targeting both may have valuable combinatorial, and perhaps even synergistic, effect.

In summary, we have shown that SLAMF6 is a constitutive inhibitory immune receptor; in its absence, CD8 T cells acquire stronger reactivity against tumor cells. The strong effector trait is attributed to a series of T-bet-mediated transcriptional events that drive CD8 T cells to exert strong cytotoxicity and achieve long-lasting tumor control. SLAMF6 is an attractive target for the development of checkpoint inhibitors for systemic treatment of cancer and for the improvement of anti-tumor cellular therapies.

## Materials and Methods

### Plasmids

pCMV3-mSLAMF6 and pCMV3-negative control vectors were purchased from SINO Biological Inc, Eschborn, Germany.

### Cells

#### Melanoma cells

Mouse melanoma B16-F10/mhgp100 cells (B16-F10 melanoma cells transduced with pMSGV1 retrovirus, which encodes a chimeric mouse gp100 with the human gp100_25-33_ sequence) were a kind gift from Ken-ichi Hanada, Surgery Branch, NCI, NIH. The cells were cultured in RPMI 1640 supplemented with 10% heat-inactivated fetal calf serum (FCS), 2 mmol/L L-glutamine, and combined antibiotics (all from ThermoFisher Scientific, Massachusetts, USA). Lines were regularly tested and were mycoplasma free.

#### Aberrant LY108 expression on melanoma cells

*The* B16-F10/mhgp100 murine melanoma cell line was transfected with pCMV3—hygromycin-mSLAMF6 using lipofectamine 2000 (ThermoFisher). Hygromycin-resistant melanoma cells were sub-cloned, and the stably transfected cells were labeled with anti-LY108 Ab and sorted in an ARIA-III sorter. B16-F10/mhgp100 cells transfected with the empty vector were sub-cloned using hygromycin resistance.

#### Tumor-infiltrating lymphocytes (TILs)

Fresh tumor specimens taken from resected metastases of melanoma patients were used to release TILs using a microculture assay (Lotem *et al.*, 2002). Human lymphocytes were cultured in complete medium (CM) consisting of RPMI 1640 supplemented with 10% heat-inactivated human AB serum, 2 mmol/l L-glutamine, 1 mmol/l sodium pyruvate, 1% nonessential amino acids, 25 mmol/l HEPES (pH 7.4), 50 μmol/l 2-ME, and combined antibiotics (all from ThermoFisher). CM was supplemented with 6000 IU/ml recombinant human IL-2 (rhIL-2, Chiron, California, USA).

#### Cloning of peptide-specific TILs

On day 14 after TIL initiation, lymphocytes were washed with PBS, re-suspended in PBS supplemented with 0.5% bovine serum albumin (BSA), and stained with FITC-conjugated HLA-A*0201/MART-1_26–35_ dextramer (Immudex, Copenhagen Denmark) for 30 min at 4°C. Lymphocytes were then incubated with allophycocyanin-conjugated mouse anti-human CD8 (eBioscience, California, USA) for an additional 30 min at 4°C and washed. CD8 lymphocytes, positively stained with the dextramer (CD8dextramer^+^ cells), were sorted with a BD Biosciences FACS Aria and directly cloned at one or two cells per well in 96-well plates in the presence of 30 ng/ml anti CD3 (eBioscience), 6000 IU/ml rhIL-2, and 4 Gy-irradiated 5×10^4^ allogeneic PBMCs as feeder cells. Five days later, 6000 IU/ml rhIL-2 was added and renewed every 2 days after that. On day 14, the clones were assayed for IFN-γ secretion in a peptide-specific manner following their co-incubation with T2 cells pulsed with MART-126–35 (commercially synthesized and purified [>95%] by reverse-phase HPLC by Biomer Technology, Cheshire, UK) by ELISA (R&D Systems, Minneapolis, MN, USA). The MART-126–35–-reactive clones were further expanded in a second-round of exposure to 30 ng/ml anti-CD3, and 6000 IU/ml rhIL-2 in the presence of 50-fold excess irradiated feeder cells.

### Antibodies

For flow cytometry, cells were labeled with the following reagents: anti-CD16/32 (93), anti-SLAMF6 (330-AJ), anti-TNFa (MP6-XT22), anti-CD19 (6D5), anti-CD44 (IM7), anti-TIM3 (RMT3-2.3), and anti-LAG-3 (C9B7W) (all from Biolegend, San Diego, CA, USA). Anti-IFN-γ (XMG1.2), anti-CD8 (53-6.7), anti-GZMB (NGZN), anti-CD4 (GK1.5), and anti-CD25 (PC61.5) were from Biogems, Westlake Village, CA, USA. Anti-CD62L (MEL-14), anti-Vb13 (MR12-3), anti-CD69 (H1.2F3), anti-CD137 (17B5), anti-PD1 (J43), and anti-CD244 (eBio244F4) were from eBioscience. Anti-human SLAMF6 (REA) was purchased from Miltenyi Biotec, Bergisch Gladbach, Germany. Anti-pS6 (D57.2.2E) was from Cell Signaling Technology, Danvers, MA, USA.

Anti-LAG-3 (C9B7W) and the corresponding isotype were from InVivoMab, BioXcell, NH, USA. The Mart-1 26-35 iTag MHC tetramer was from MBL, Woburn, MA, USA.

For Immunobloting: anti-Ly108 (Rat, 3E11, Merck, Kenilworth, NJ, USA), anti-SAP (Rat, 1A9, Biolegend), anti-β actin (Mouse, sc-47778, Santa Cruz Biotechnology, Texas, USA), anti-SHP1 (Rabbit, generated in A.V.’s laboratory).

### Mice

C57BL/6 mice were purchased from Harlan laboratories. Pmel-1 (a kind gift from M. Baniyash) and SLAMF6 KO mice (a kind gift from I. Shachar) were self-bred.

The Pmel-1 mice carry a rearranged TCR specific for a 9-mer epitope (25-32) from murine Pmel 17, overexpressed on transformed melanocytes and homologous to the human melanoma-associated antigen gp100.

All experiments were performed with 8- to 12-week-old female mice.

#### Generation of Pmel-1 x SLAMF6 KO mice

Pmel-1 and SLAMF6−/− mice were bred to generate Pmel-1 X SLAMF6−/− mice according to the ethics requirements (Authority for biological and biomedical models, Hebrew University, Jerusalem, Israel).

When the mice reached 3 weeks of age, 2 mm of the mouse tail were cut, 200 μl 50 mM NaOH 0.2Mm EDTA were added, and the tails were incubated at 95°C for 20 min for DNA purification. 200 μl 80 mM TRIS-HCL, pH5, were added to stop the reaction. The DNA purified from the tails was used in PCR reactions for genotyping of mice in the SLAMF6 locus on chromosome 1 (primers adapted from the Jackson laboratories website) and in the Pmel-1 locus on chromosome 2 (Ji *et al.*, 2014). The identification of the genomic insertion site of the Pmel-1 TCR α and β transgenes was performed by next-generation sequencing.

### Splenocyte activation

Pmel-1 or Pmel-1xSLAMF6−/− mouse splenocytes (2×10^6^/ml) were activated with 1μg/ml of mouse gp100_25-33_ peptide for 6 days with IL-2 30 IU/ml. Fresh medium containing IL-2 was added every other day.

### *In vitro* assays

#### RNA isolation and qPCR

RNA was isolated from cells using the GenElute Mammalian Total RNA kit (Sigma Aldrich, MA, USA) according to the manufacturer’s protocol. RNA was then transcribed to cDNA using qScript cDNA Synthesis kit (Quantabio, Beverly, MA, USA) according to the manufacturer’s instructions, and RT-PCT or qRT-PCR was performed using the following primers:

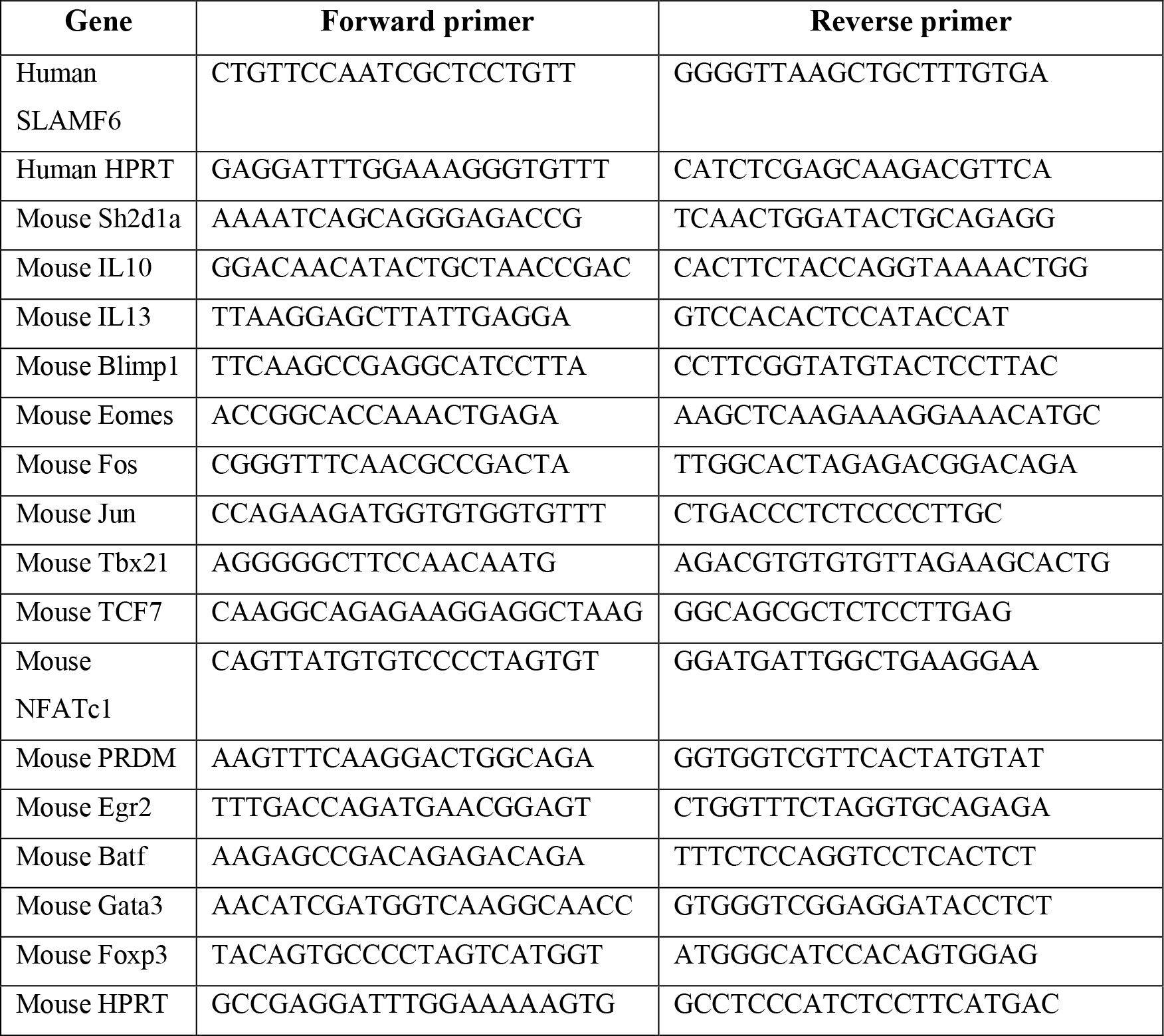

RT-PCR for *Sh2d1a* was performed in the SensQuest lab cycler machine (Danyel Biotech); the products were then run on 1.5% agarose gel.

#### Longitudinal expression of SLAMF6

TIL412 was activated using plate-bound anti-CD3 1μg/ml supplemented with IL-2. On every other day, 1×10^5^ cells were stained in triplicates to evaluate SLAMF6 expression using anti-SLAMF6 (Miltenyi Biotec) or the corresponding isotype control. Pmel-1 splenocytes were activated using 1μg/ml mouse gp100_25-33_ peptide supplemented with 30 IU/ml IL-2 for 6 days, and SLAMF6 expression was tested at the indicated time points using anti-SLAMF6 (Biolegend, clone 330-AJ).

#### RNA sequencing of activated CD4 cells

Human PBMCs were isolated by Ficoll-Paque (GE Healthcare Life Sciences, Pittsburgh, PA, USA) centrifugation from blood drawn from donors recruited from the Boston community as part of the Phenogenetic Project and ImmVar Consortium. Naive CD4 T cells were isolated from PBMCs by negative selection using the human Naive CD4^+^ T Cell Isolation Kit II (Miltenyi Biotec). The isolation was >97% pure as assessed by flow cytometry. For each condition, 50,000 T cells were used. Beads used for cell stimulation were generated by incubating 4×10^6^ CELLection Pan Mouse IgG beads (ThermoFisher) with antibodies (BD Biosciences, San Jose, CA, USA) against CD3 (UCHT1; 10.67 ng) and CD28 (CD28.2; 5.33 ng) complemented to 20 ng total protein with control IgG1 (MOPC-31C) for CD3/CD28 stimulation at the following time points: 0, 3, 6, 12, 24, and 72 hours. RNA for sequencing was isolated using an RNeasy 96 kit (Qiagen, Hilden, Germany), treated with DNase I (New England Biolabs, Ipswich, MA, USA), and converted to sequencing libraries using the Smartseq2 protocol (PMID: 24385147). Libraries were sequenced on a Novaseq S2 (Illumina, San Diego, CA, USA) with paired-end 50bp reads.

#### RNA sequencing results analysis

We used Bowtie v1.0.0 to align raw data to the UCSC hg19 transcriptome and RSEM v1.2.8 to quantify gene expression levels (TPM – transcripts per million). Immune-related genes were selected from the RNA sequencing results based on a unified list of genes created from the Immunogenetic Related Information Source (IRIS) list and the MAPK/NFKB Network list (https://www.innatedb.com/redirect.do?go=resourcesGeneLists).

For the immune-related genes, heatmaps were generated using the Morpheus online tool (https://software.broadinstitute.org/morpheus/).

#### Intracellular cytokine staining

Mouse splenocytes (1×10^5^) were co-cultured for 6h at 37°C at a 1:1 ratio with the indicated target melanoma cells or activated with 1μg/ml anti-CD3 (Biolegend, clone: 145-2C11). After 2 h, Brefeldin A (Biolegend) was added (1:1000). After incubation the cells were washed twice with PBS and stained with anti-CD8 (Biogems) for 30 min at room temperature. Following fixation and permeabilization (eBioscience protocol), intracellular IFN-γ and TNF-α were labeled with anti-IFN-γ and anti-TNF-α antibodies, respectively (TNF-α, clone: MP6-XT22, Biolegend, and IFN-γ, clone: XMG1.2, Biogems) for 30 min at room temperature. Cells were washed with permeabilization buffer, resuspended in FACS buffer, and subjected to flow cytometry.

#### Intracellular staining for phosphorylated proteins

For intracellular staining of phosphorylated proteins, cells were activated for different times, fixed with lyse/fix buffer (BD Biosciences), permeabilized with Perm II buffer (BD Biosciences) and stained with fluorescence-labeled antibodies against pS6 (Cell Signaling, D57.2.2E) in stain buffer (BD Biosciences).

#### Cell viability assay

Following expansion, splenocytes were washed, and 1×10^5^ cells were cultured in CM. After 7 days, cells were washed and labeled with the Annexin V apoptosis detection kit (eBioscience), according to the manufacturer’s instructions. Cells were analyzed by flow cytometry.

#### Proliferation assay

Fresh splenocytes were labeled with CFSE (Quah, Warren and Parish, 2007) and activated as described above. At the indicated days, cells were counted, labeled with anti-CD8 Ab (Biogems, clone: 53-6.7), and subjected to flow cytometry.

#### Interferon-gamma secretion

Mouse splenocytes, previously activated for 7 days, were co-cultured (1×10^5^) overnight at a 1:1 ratio with the indicated target melanoma cells or activated with 1μg/ml plate-bound anti - CD3 (Biolegend, clone: 145-2C11) as indicated in each experiment. Conditioned medium was collected, and mouse IFN-γ secretion was detected by ELISA (Biolegend).

#### GM-CSF secretion

Mouse splenocytes, previously activated for 7 days, were co-cultured (1×10^5^) overnight at a 1:1 ratio with the indicated target melanoma cells. Conditioned medium was collected, and mouse GM-CSF secretion was measured by ELISA (Biolegend).

#### Immunohistochemistry

For histological analysis, spleen and tumor tissues were cut into 5 μm sections, deparaffinized with xylene, and hydrated with graded ethanol. Endogenous peroxidase was blocked using 3% H_2_O_2_ for 5 min. and 0.01M citrate buffer (pH 6.0) was used for antigen retrieval. The samples were cooked in a pressure cooker at maximum power for 13 min and then at 40% power for an additional 15 min, and left to cool for 30 min until they reached room temperature. All slides were then washed in PBS and blocked for 30 min with CAS block at room temperature. The tissues were incubated with rabbit anti-CD4 Ab (ab183685) or rabbit anti-CD8 Ab (ab203035) diluted in CAS-Block (1:1000 or 1:250, respectively) overnight at 4°C. The following day, the slides were washed in PBS, incubated with Dako anti-rabbit secondary antibody for 30 min and developed with AEC for 10 min. After several washes in PBS and dH_2_O, the slides were counterstained with hematoxylin, rinsed in H_2_O, and covered with fluoromount (ThermoFisher) and a cover-slip. The photos were taken with an Olympus BX50 microscope and Olympus DP73 camera at room temperature. The acquisition software used was CellSens Entry 1.8.

#### Killing assay

Splenocytes were co-cultured for 19h at 37°C at a 1:1 ratio with the indicated target melanoma cells. After 16h, Brefeldin A (Biolegend) was added (1:1000). At the end of the incubation, the cells were washed twice with PBS and stained with aCD8 (Biogems, clone 53-6.7) for 30 min at room temperature. Following fixation and permeabilization (eBioscience protocol), intracellular granzyme-B was labeled with anti-GzmB antibody (Biogems, clone: NGZB) for 30 min at RT. Cells were washed with permeabilization buffer, resuspended in FACS buffer, and subjected to flow cytometry.

#### Cytokine array

Day 7 activated mouse splenocytes were co-cultured (1×10^5^) overnight at a 2:1 ratio with the B16-F10/mhgp100 melanoma cells. Conditioned medium was collected and used in the Quantibody mouse cytokine array (QAM-CYT-1, RayBiotech, Peachtree Corners, GA, USA).

#### Immunobloting

Cells were lysed using RIPA buffer, and protein concentrations were tested using Bradford quantification. Equal concentrations of lysates were resuspended in SDS sample buffer (250 mM Tris-HCl [pH 6.8], 5% w/v SDS, 50% glycerol, and 0.06% w/v bromophenol blue) for 5 min at 95°C. Proteins were separated by SDS PAGE and transferred to a PVDF membrane. Membranes, blocked with 1% milk solution, were incubated with primary antibodies overnight at 4°C, followed by incubation with HRP-conjugated secondary antibodies for 1 hour at RT (Jackson ImmunoResearch Laboratories). Signals were detected by enhanced chemiluminescence reagents (Clarity Western ECL Substrate, BioRad, Hercules, CA, USA).

#### Flow cytometry

After blocking Fc receptors with anti-CD16/CD32 antibody, cells were stained with antibodies or tetramers on ice or at room temperature for 25 min, according to the manufacturer’s instructions. Subsequently, cells were washed and analyzed using CytoFlex (Beckman Coulter, CA, USA), and flow cytometry-based sorting was done in an ARIA-III sorter. Flow cytometry analysis was done using FCS express 5 flow research edition (De Novo software). The gating strategy is illustrated in Supplementary Figure 1.

### *In vivo* assays

#### Adoptive cell transfer experiments

a. B16-F10/mhgp100-empty vector or B16-F10/mhgp100-SLAMF6 transfected mouse melanoma cells (0.5×10^6^) were injected s.c. into the back of C57BL/6 mice. Pmel-1 mouse splenocytes (2×10^6^/ml) were expanded with 1μg/ml of mouse gp100_25-33_ peptide in the presence of IL-2 (30 IU/ml). Fresh medium containing IL-2 was added every other day. On day 7, 10^7^ Pmel-1 cells were adoptively transferred i.v. into 500 CGy-irradiated tumor-bearing mice. 0.25×10^6^ IU/100μl IL-2 was administered i.p. twice daily for 2 days. Tumor size and mouse weight were measured 3 times a week. Follow-up was conducted until the ethical humane endpoint was reached.
b. B16-F10/mhgp100 mouse melanoma cells (0.5×10^6^) were injected s.c. into the back of C57BL/6 mice. Pmel-1 or Pmel-1xSLAMF6 KO mouse splenocytes (2×10^6^/ml) were expanded with 1μg/ml of gp100_25-33_ peptide in the presence of IL-2 (30 IU/ml). Fresh medium containing IL-2 was added every other day. On day 7, 10^7^ Pmel-1 cells or Pmel-1xSLAMF6 KO cells were adoptively transferred i.v. into 500 CGy-irradiated tumor-bearing mice. 0.25×10^6^ IU/100μl IL-2 was administered i.p. twice a day for 2 days. Tumor size and mouse weight were measured 3 times a week. Follow up was conducted until the ethical humane endpoint was reached or until day 80 (for mice that showed complete tumor remission).
c. The experiment was performed as in (b). One week after ACT, mice were sacrificed, and tumors, DLN, and spleens were harvested for further analysis by flow cytometry or immunohistochemistry.

Animal studies were approved by the Institutional Review Board - Authority for biological and biomedical models, Hebrew University, Jerusalem, Israel (MD-14602-5 and MD-15421-5).

#### Statistics

Statistical significance was determined by unpaired t-test (two-tailed with equal SD) using Prism software (GraphPad). A p-value < 0.05 was considered statistically significant. Analysis of more than two groups was performed using one-way ANOVA test. *, p≤0.05; **, p≤0.01; ***, p≤0.001. For each experiment, the number of replicates performed and the statistical test used are stated in the corresponding figure legend.

## Acknowledgments

The authors wish to acknowledge the devoted technical work of Anna Kuznetz and Yael Gelfand. We thank Eli Pikarsky and Ofer Mandelboim for helpful discussions, and Karen Pepper for editing the manuscript.

We thank Kathleen B. Yates and W. Nicholas Haining, who provided training and their expertise towards the design of a range of beads to optimize control of the degree and nature of antigenic stimulation of T cells *in vitro*.

---

**Supplementary Figure S1:**
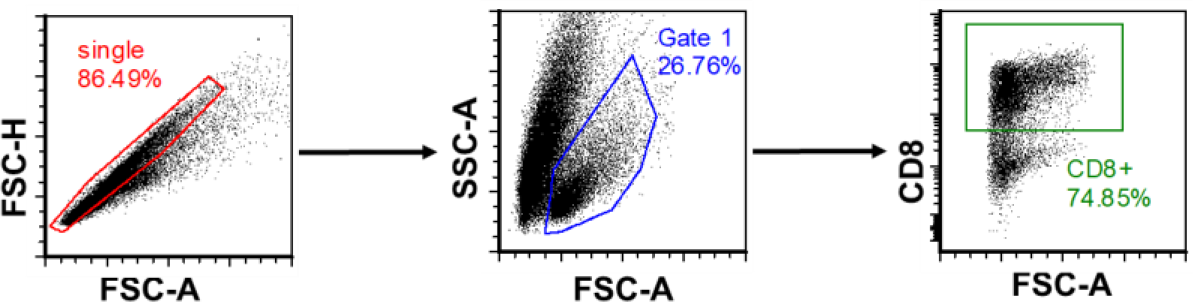
Gating strategy. Initially, non-single cells were excluded using FSC-A and FSC-H axes. Then, based on the morphology of the cells in the FSC-A SSC-A axes, the live cell population was gated. In this population, only cells that stained positively for CD8 expression were subjected to further analysis.

**Supplementary Figure S2:**
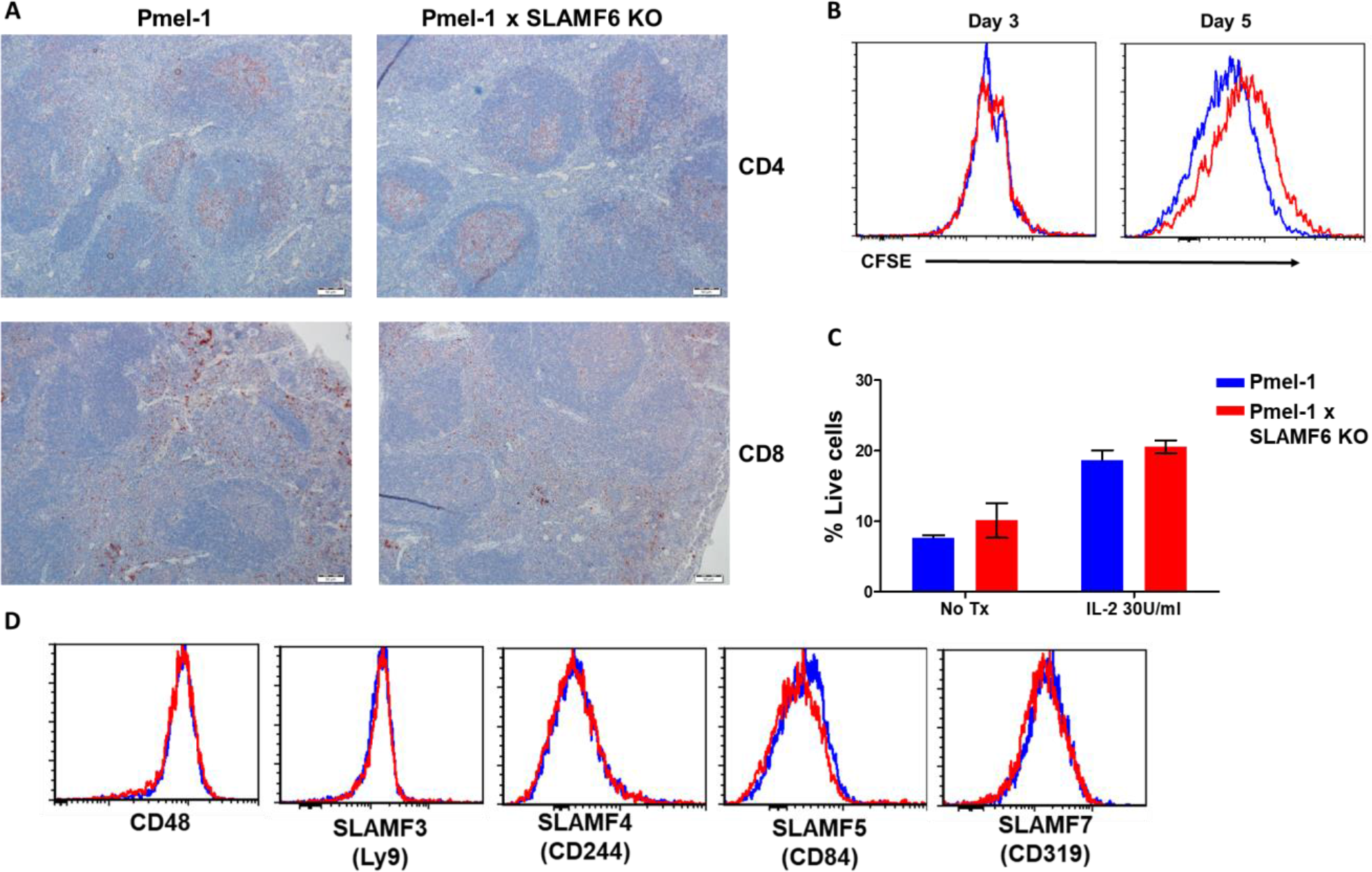
**A**, Immunohistochemistry staining of Pmel-1 and Pmel-1 x SLAMF6 KO spleen sections using anti CD4 and anti-CD8 antibodies (X10 magnification). **B**, Pmel-1 and Pmel-1 x SLAMF6 KO splenocytes were labeled with CFSE and activated; at the indicated time points the cells were stained for CD8 expression, and CFSE level gated on the CD8 population was measured using flow cytometry. **C**, Pmel-1 and Pmel-1 x SLAMF6 KO splenocytes activated for seven days followed by seven days maintenance with IL-2 (30 IU/ml) or without its addition. Percentage apoptotic and dead cells was measured by PI-Annexin V. Summary of two experiments shown. No Tx, no treatment. **D**, After 7 days of activation, Pmel-1 and Pmel-1 x SLAMF6 KO CD8 T cells were stained with antibodies against SLAM family receptors. The expression level of each receptor in CD8 cells is presented.

**Supplementary Figure S3:**
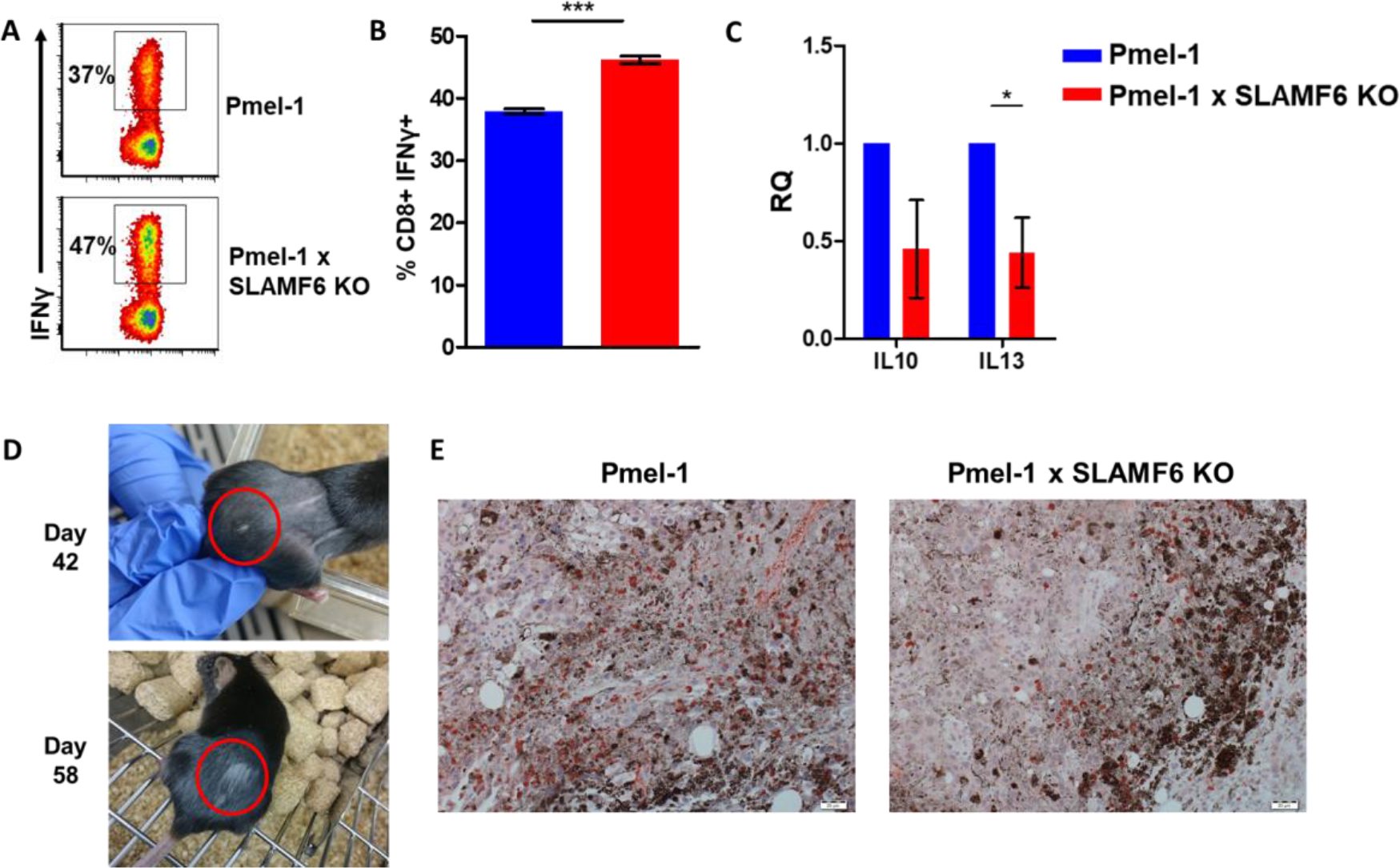
**A, B**, Pmel-1 or Pmel-1 x SLAMF6 KO splenocytes were activated for 7 days with gp100_25-33_ peptide and IL-2 (30 IU/ml). Cells were then incubated for 6 hours with B16-F10/mhgp100 melanoma cells. IFN-γ production was detected by flow cytometry. One representative experiment (**A)**, and a summary of triplicates (**B)**, are shown. **C**, Pmel-1 and Pmel-1 x SLAMF6 KO splenocytes were activated for 7 days and lysed. RNA was extracted and quantitative RT-PCR for cytokine expression was performed. Data were normalized to *Hprt* expression for each mouse strain. Pmel-1 x SLAMF6 KO values for each gene were normalized to Pmel-1 values. **D**, Photographs from days 42 and 58 post-tumor inoculation of a mouse that developed vitiligo following ACT with Pmel-1 x SLAMF6 KO cells. Vitiligo spots are marked. **E**, Immunohistochemistry staining of tumors from mice receiving ACT of Pmel-1 or Pmel-1 x SLAMF6 KO splenocytes, harvested 7 days post ACT. Tumor sections were stained with anti-CD8 Ab (X20 magnification).

**Supplementary Figure S4:**
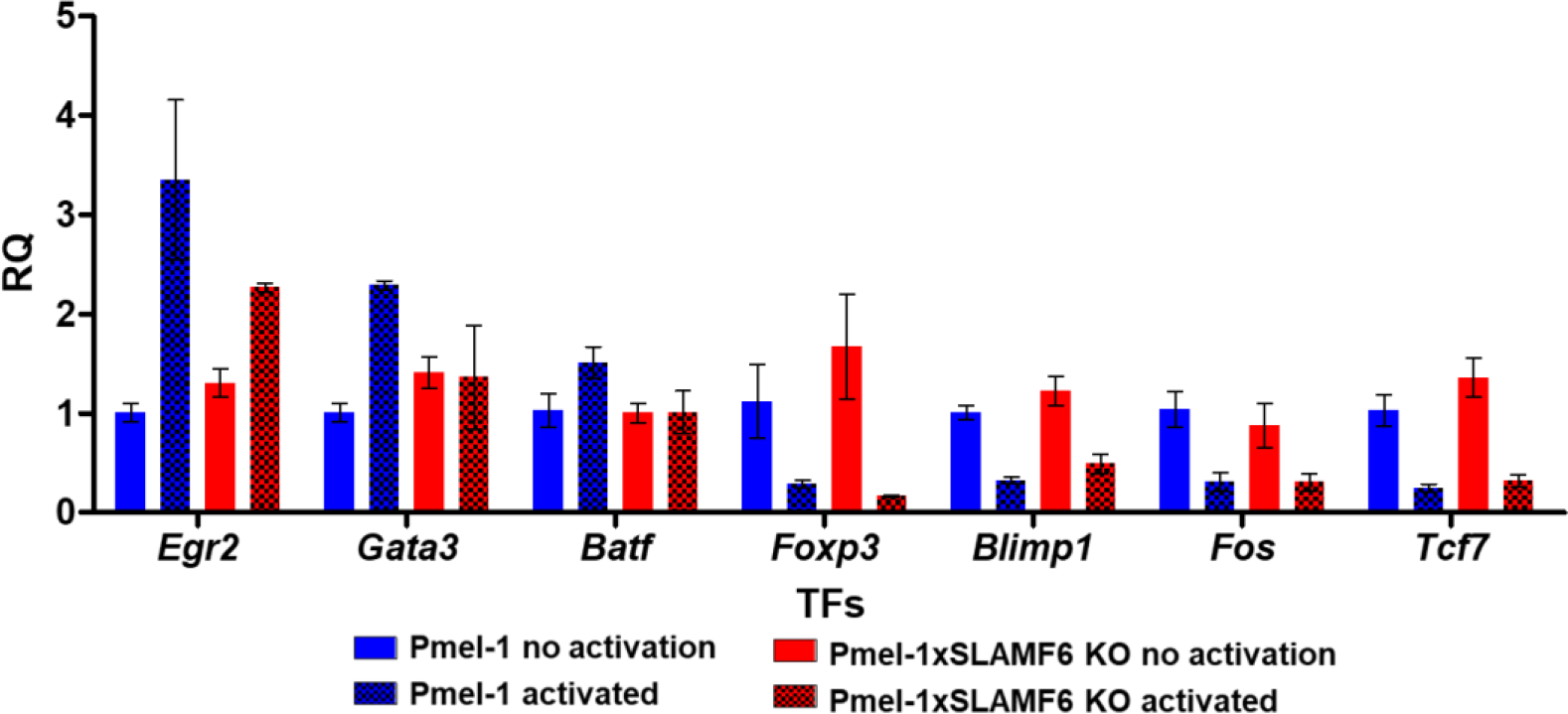
Pmel-1 and Pmel-1 x SLAMF6 KO splenocytes were either activated with gp100_25-33_ peptide in the presence of IL-2 (30 IU/ml) for 18 hours or only kept with IL-2 (non-activated). After 18 hours, the cells were lysed, RNA was extracted, and quantitative RT-PCR for transcription factors expression was performed. Data was normalized to *Hprt* expression for each mouse strain. Values for each condition were normalized to Pmel-1 non-activated values for each gene.

